# *Gata2*-regulated *Gfi1b* expression controls endothelial programming during endothelial-to-hematopoietic transition

**DOI:** 10.1101/2022.03.24.485642

**Authors:** Cansu Koyunlar, Emanuele Gioacchino, Disha Vadgama, Hans de Looper, Joke Zink, Remco Hoogenboezem, Marije Havermans, Eric Bindels, Elaine Dzierzak, Ivo P Touw, Emma de Pater

## Abstract

The first hematopoietic stem cells (HSCs) are formed through endothelial-to-hematopoietic transition (EHT) events during embryonic development. The transcription factor *GATA2* is a crucial regulator of EHT and HSC function throughout life. Because *GATA2* haploinsufficiency patients have inborn mutations, prenatal defects are likely to have an influence on disease development. In mice, *Gata2* haploinsufficiency (*Gata2*^*+/-*^) reduces the number and the functionality of embryonic hematopoietic stem and progenitor cells (HSPCs) generated through EHT. However, the embryonic HSPC pool is heterogeneous and the mechanisms underlying this defect in *Gata2*^*+/-*^ embryos are unclear. Here, we investigated whether *Gata2* haploinsufficiency selectively affects a cellular subset undergoing EHT. We show that *Gata2*^*+/-*^ HSPCs initiate but cannot fully activate hematopoietic programming during EHT. In addition, due to reduced activity of the endothelial repressor *Gfi1b, Gata2*^*+/-*^ HSPCs cannot repress the endothelial identity to complete maturation. Finally, we show that hematopoietic-specific induction of *gfi1b* can restore HSC production in *gata2b-*null (*gata2b*^*-/-*^) zebrafish embryos. This study illustrates pivotal roles of *Gata2* on the regulation of transcriptional network governing HSPC identity throughout EHT.

**Highlights:**

- Maturation of embryonic *Gata2*^*+/-*^ HSPCs is disturbed due to aberrant endothelial gene expression and incomplete activation of hematopoietic transcriptional programming.
- *Gata2* activates *Gfi1b* to repress endothelial identity of embryonic HSPCs during maturation.
- Hematopoietic-specific induction of *gfi1b* restores the number of embryonic HSCs in *gata2b*^*-/-*^ zebrafish.

## Introduction

Hematopoiesis relies on multipotent, self-renewing HSCs. HSCs originate from the ventral wall of the embryonic dorsal aorta at the aorta-gonad-mesonephros (AGM) region. In the AGM region, definitive HSPCs are generated through a trans-differentiation process from a specialized endothelial cell (EC) compartment with hematopoietic potential (hemogenic endothelial cells or HECs). This process of endothelial-to-hematopoietic transition (EHT) is conserved between mammalian and non-mammalian vertebrates *(Ciau-Uitz and Patient, 2019; Ivanovs et al., 2017; Bertrand et al., 2010; Boisset et al., 2010; Kissa and Herbomel, 2010; Zovein et al., 2008; de Bruijn et al., 2002; North et al., 2002; Medvinsky and Dzierzak, 1996; Muller et al., 1994; Dieterlen-Lievre, 1975)*. In mice, EHT events occur between embryonic days (E)10.5-E12.5. Phenotypic HSPCs emerge from intra-aortic hematopoietic clusters (IAHCs) through EHT and co-express endothelial markers such as CD31 and hematopoietic markers like cKit *(Yokomizo and Dzierzak, 2010)*. Previous studies showed that HSPCs develop through a multistep maturation process within IAHCs and that only a small fraction of IAHC cells become multipotent HSCs *(Rybtsov et al., 2014; 2011; Taoudi et al., 2008)*. Throughout AGM maturation, HSPCs gradually repress the endothelial-specific gene expression and upregulate the expression of hematopoietic-specific genes *(Oatley et al., 2020; Baron et al., 2018; Zhou et al., 2016; Swiers et al., 2013)*. Following their emergence from the AGM, HSCs migrate to the fetal liver and eventually colonize the BM around birth to maintain the hematopoietic system throughout life *(Zovein et al., 2008)*.

The transcription factor *GATA2* is one of the key regulators of hematopoietic programming. In mice, *Gata2* is required for embryonic HSC generation and survival. While germline deletion of *Gata2* (*Gata2*^*-/-*^) is lethal at E10, i.e., just before the appearance of the first HSCs, *Gata2* haploinsufficiency (*Gata2*^*+/-*^) severely reduces the number of embryonic HSPCs, but these heterozygous mice survive to adulthood despite reduced numbers of HSCs *(Gao et al., 2013; de Pater et al., 2013; Rodrigues et al., 2005; Ling et al., 2004; Tsai et al., 1994)*. Moreover, conditional deletion of *Gata2* in HECs does not fully abrogate the formation of IAHCs, but depletes the functional HSCs *(de Pater et al., 2013)*. In addition, HSCs do not survive from conditional deletion of *Gata2* after their emergence and become apoptotic *(de Pater et al., 2013)*.

Clinical manifestation of germline heterozygous *GATA2* mutations results in *GATA2* haploinsufficiency syndromes in patients. Typically, patients present with bone marrow failure and are at a high (80%) risk of developing myelodysplastic syndrome (MDS) or acute myeloid leukemia (AML) before the age of 40 *(Donadieu et al., 2018; Spinner et al., 2014; Dickinson et al., 2011; Hsu et al., 2011; Ostergaard et al., 2011; Hahn et al., 2011)*. Although *GATA2* expression is required in HSCs during both embryonic and adult stages, the consequences of embryonic *GATA2* haploinsufficiency for disease development are still unexplored.

Despite the requirement for *Gata2* during EHT, mechanisms specifically involved in the production and growth of HPSCs in healthy and GATA2 haploinsufficiency states are incompletely understood. Here we sought to understand why *GATA2* haploinsufficiency depletes phenotypic HSCs. In the present study, we explore how embryonic *Gata2* haploinsufficiency affects EHT and the development of first phenotypic HSCs in the mouse AGM. We show that hematopoietic programming was not abrogated in *Gata2*^*+/-*^ E11 HSPCs. However, maturation of *Gata2*^*+/-*^ HSPCs was disturbed and transcriptional profiling showed that *Gata2* is the key factor in the gene regulatory network in nascent HSCs. Importantly we demonstrate that *Gata2* regulates *Gfi1b* to repress endothelial gene expression during HSPC maturation. Ectopic expression of *gfi1b* restores the number of phenotypic HSCs in *gata2b*-deficient zebrafish embryos to reveal this previously unidentified role of *Gata2* on modulating the transcriptional programming and maturation of HSPCs during EHT.

## Materials and Methods

### Mouse and zebrafish models

*Gata2*^*+/-*^ mice (*Tsai et al., 1994*), *gata2b*^*-/-*^ zebrafish (*Gioacchino et al., 2021*) and *Tg(CD41:GFP)* zebrafish (*Ma et al., 2011*) were previously described. All animals were housed and bred in animal facilities at the Erasmus MC, Rotterdam, Netherlands. Animal studies were approved by the Animal Welfare and Ethics Committees of the EDC in accordance with legislation in the Netherlands.

### Whole-mount immunofluorescence staining

E11 WT and *Gata2*^*+/-*^ AGMs were dissected, prepared and mounted as previously described (*Yokomizo et al., 2012*). Primary antibody CD117 (cKit) rat anti-mouse (Invitrogen) was combined with secondary antibody Alexa Fluor-488 goat anti-rat (Invitrogen) for cKit visualization. CD31 was visualized by using biotinylated CD31 rat anti-mouse (BD Biosciences) and Cy5-conjugated Streptavidin (Jackson Immunoresearch) primary and secondary antibodies respectively. Whole AGM region for each sample was imaged using Leica SP5 confocal microscope. CD31 and cKit double positive cells were analyzed using Leica Application Suite X (version 4.3) software.

### RNA isolation and sequencing

Cells were sorted in Trizol (Sigma) and total RNA isolation was performed according to the standard protocol using GenElute LPA (Sigma). RNA quality and quantity was assessed on 2100 Bioanalyzer (Agilent) using RNA 6000 Pico Kit (Agilent). cDNA was prepared using SMARTer procedure with SMARTer Ultra Low RNA kit (Clontech) and sequenced on Novaseq 6000 platform (Illumina).

### Gene set enrichment and network analysis

Gene expression values were measured as FPKM (Fragments per kilobase of exon per million fragments mapped) and differential expression analysis was performed using the DESeq2 package in the R environment. Gene set enrichment analysis (GSEA) was performed on the FPKM values using the curated gene sets in the Molecular Signatures Database (MSigDB). GSEA results were used as an input for network analysis performed in Cytoscape software.

### ATAC sequencing

Cells were processed for library preparation using the previously described protocol by Delwel group (*Ottema et al., 2021*). Libraries were quantified using Qubit and NEBNext Library Quant Kit for Illumina (NEB). Quality of the libraries were determined by the peak distribution visualization using 2100 Bioanalyzer (Agilent). Samples were sequenced on Novaseq 6000 platform (Illumina). Bigwig files were generated using the bamCoverage tool from deepTools and visualized using Integrative Genomics Viewer (IGV) software.

### Flow cytometry and sorting (FACS)

AGM regions were dissected as described before (*Yokomizo et al., 2012*). Tissues were incubated with collagenase I (Sigma) in phosphate buffer solution (PBS) supplemented with 5 IU/mL penicillin, 5 μg/mL streptomycin, and 10% fetal calf serum (FCS) for 45 min at 37°C. Cells were stained using antibodies: PE-Cy7 anti-mouse CD31 (eBioscience), APC rat anti-mouse CD117 (cKit, BD Bioscience), FITC anti-mouse CD41 (Biolegend), PE rat anti-mouse CD43 (BD Bioscience) and Alexa Fluor-700 rat anti-mouse CD45 (BD Bioscience). All antibody incubations were performed in PBS + 10% FCS for 30 min on ice. After washing with PBS + 10% FCS at 1000 rpm for 10 min, cell pellets were resuspended with 1:1000 DAPI in PBS + 10% FCS for live/death cell discrimination. FACS events were recorded and cells were sorted using FACSAria III (BD Biosciences). Results were analyzed and visualized using FlowJo 7.6.5 software.

### Colony-forming unit assay

Cells were incubated in MethoCult GF M3434 (Stem Cell Technologies) supplemented with 5 IU/mL penicillin and 5 μg/mL streptomycin at 37°C. Colony-forming units (CFU) were scored after 11 days of culture. Growth of primitive erythroid progenitor cells (BFU-E) and granulocyte-macrophage progenitor cells (CFU-GM, CFU-G and CFU-M) were scored using an inverted microscope.

### Generation of a *gfi1b* construct

Wild type sequences of *gfi1b, mCherry* and *runx1* +23 enhancer were separately cloned into pJET1.2 vectors using CloneJet PCR Cloning Kit (Thermo Fisher). Following transformation, outgrown colonies were picked for DNA isolation. Presence of the insert was confirmed by restriction enzyme digestion using BglII (NEB) followed by agarose gel electrophoresis (1,5 %). DNA fragments then used as a PCR template for the Gibson cloning reaction. Fragments were amplified using overhang primers (Supplementary table 1) and purified using DNA Clean & Concentrator kit (Zymo Research). The pUC19-iTol2 backbone was digested with BamHI-HF (NEB) overnight at 37°C and NEBuilder HiFi Assembly MasterMix (NEB) was used for the assembly of the fragments. Correct assembly was determined by HindIII restriction enzyme digestion and PCR amplification for the fragments. All transformations were done using E. coli and by performing heat shock for 30 seconds at 42°C followed by recovery in SOC outgrowth medium (NEB) for 1 hour. All colonies were grown in LB medium plates supplemented with carbenicillin (50 mg/ml; 1000:1 v/v) and DNA from individual colonies were isolated using QIAprep Spin Miniprep Kit (Qiagen).

### Generation of mRNA transposase

Plasmid with iTol2 sequence linearized using Not1 restriction enzyme (NEB). Linearized DNA was used as a template and RNA synthesis was performed using the HiScribe SP6 RNA Synthesis Kit (NEB) according to manufacturer’s instructions. mRNA was precipitated using 3M sodium acetate (1:100) and 100% ethanol (3:1) and incubated overnight at -20 °C. Transposase mRNA was verified using 0.7% agarose gel electrophoresis.

### Microinjection and embryo selection

Injection needles were prepared using the P-30 Magnetic Glass Microelectrode Vertical Needle Puller (Sutter Instrument). *gfi1b* construct and mRNA transposase were co-injected to the single cell of WT(*CD41:GFP*) or *gata2b*^*-/-*^(*CD41:GFP*) zebrafish embryos at 1-cell stage using PV830 Pneumatic PicoPump (WPI). Embryos were anesthetized using 160 mg/L Tricaine (Sigma) for the selection of reporter expression. Reporter expression was assessed using the Leica DMLB fluorescence microscope.

### *cmyb* in situ hybridization

Following injections, embryos were treated with 0.003% 1-phenyl-2-thiourea (PTU, Sigma) at 24 hpf and fixed overnight with 4 % paraformaldehyde (PFA) in phosphate-buffered saline (PBS) containing 3% sucrose at 33 hpf. In situ hybridization (ISH) for *cmyb* was performed as previously described *(Gioacchino at al., 2021; Chocron et al., 2007)*. The *cmyb* probe was a gift from Roger Patient. Results were imaged using an inverted microscope.

## Statistics

Statistical analysis was carried out in GraphPad Prism 8.0.1 software. Normally distributed data were analyzed using unpaired t-test and otherwise Mann-Whitney test was used. Significance cut-off was set at P < 0.05.

## Results

### E11 Gata2^+/-^ HSPCs undergo incomplete EHT

To explore the mechanisms leading to diminished HSPCs in *Gata2*^*+/-*^ embryos, we sorted CD31^+^cKit^+^ HSPCs from E11 WT and *Gata2*^*+/-*^ AGMs and performed RNA-sequencing (RNA-seq) experiments (Figure 1A). We found significant differences in principal component analysis (PCA) between the transcriptomic signatures of WT and *Gata2*^*+/-*^ HSPCs (Figure 1B). To understand the biological processes affected by the differentially expressed genes between the two genotypes, we performed gene-set enrichment analysis (GSEA) using curated gene-sets and compared our data to a previously published dataset showing upregulated and downregulated gene-sets found in EC, HEC and HSPC compartments during EHT (*Solaimani et al., 2015*). A hematopoietic-specific gene-set *Hematopoiesis stem cell* (upregulated in HECs and HSPCs compared to ECs) was overrepresented in *Gata2*^*+/-*^ HSPCs compared to WT, indicating that there was no defect in the initiation of hematopoietic programming in these cells (Figure 1C). Surprisingly, an endothelial-specific gene-set *Vegfa Targets*, which was downregulated in HSPCs compared to ECs and HECs, was also enriched in *Gata2*^*+/-*^ HSPCs (Figure 1D). The upregulation of both hematopoietic and endothelial signatures *Gata2*^*+/-*^ HSPCs suggests that these cells can initiate but are not able to complete EHT.

**Figure 1.**
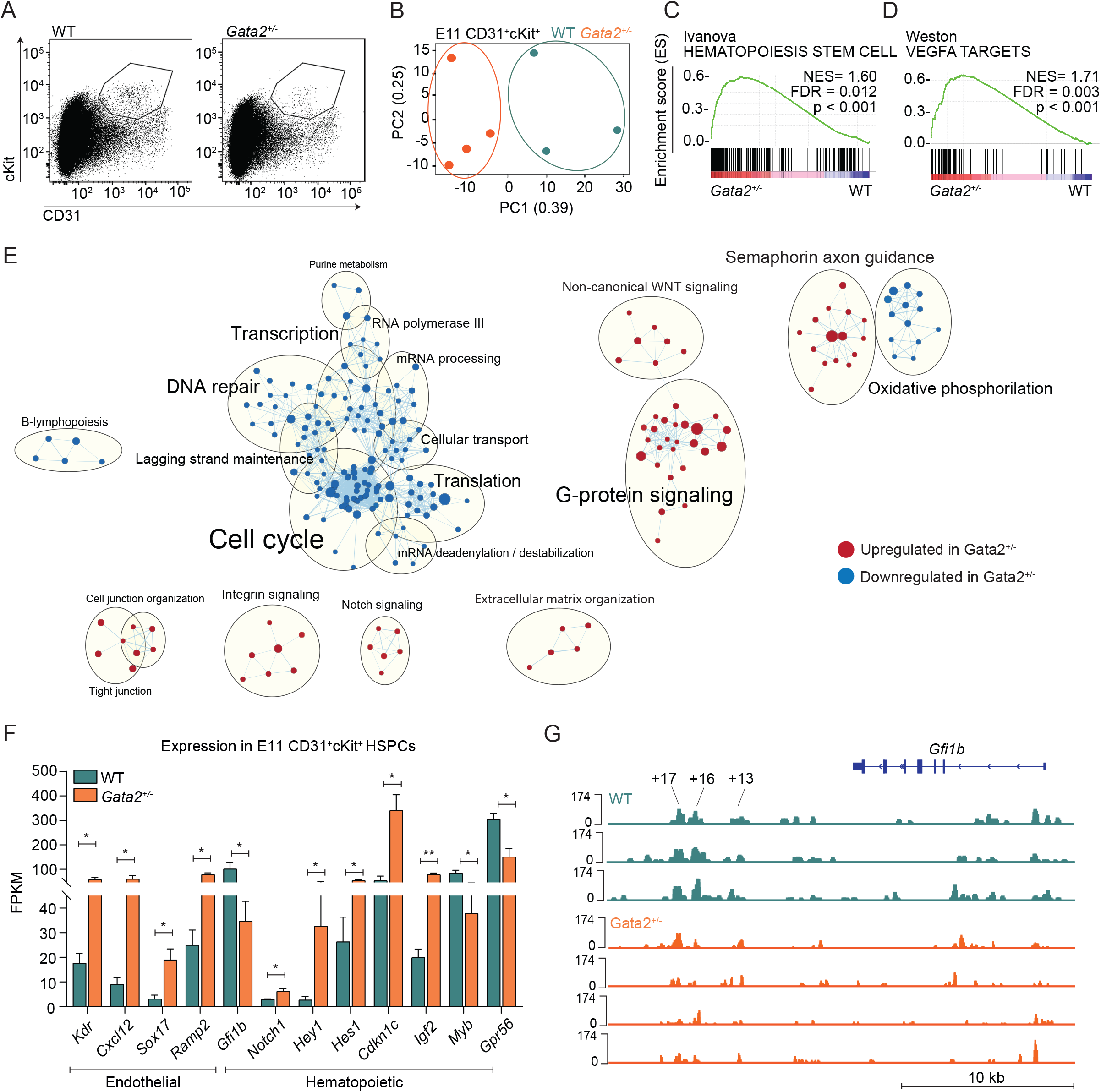
E11 *Gata2*^*+/-*^ HSPCs have aberrant hematopoietic and endothelial transcriptome. A) Sorting strategy for CD31^+^cKit^+^ cells from E11 WT (left) or *Gata2*^*+/-*^ (right) embryos. B) PCA of E11 WT (green) and *Gata2*^*+/-*^ (orange) HSPCs. Dots represent the transcriptome of CD31^+^cKit^+^ cells from individual embryos (WT; n=4, *Gata2*^*+/-*^ n=3). C-D) Gene sets upregulated in *Gata2*^*+/-*^ HSPCs compared to WT HSPCs in GSEA; C) *Hematopoiesis stem cell* and D) *Vegfa targets*. E) Network analysis comparing E11 WT and *Gata2*^*+/-*^ HSPCs. Red dots show upregulated and blue dots show downregulated gene sets in *Gata2*^*+/-*^ HSPCs compared to WT. F) Comparison of the FPKM values of endothelial (*Kdr, Cxcl12, Sox17, Ramp2*) and hematopoietic (*Gfi1b, Notch1, Hey1, Hes1, Cdkn1c, Igf2, Myb, Gpr56*) specific genes between WT and *Gata2*^*+/-*^ HSPCs. G) Comparison of open chromatin between CD31^+^cKit^+^ cells isolated from individual E11 WT (N=3, green) or *Gata2*^*+/-*^ (N=4, orange) embryos visualized by using IGV software. Accessible chromatin for *Gfi1b* and its + 13, + 16 and + 17 distal enhancer regions were analyzed. Peak range is set to min=0 and max= 174 for all samples. Tool bar represents 10 kb in length. Error bars represent standard error of the mean. *P < 0.05, **P < 0.01.

To further investigate the transcriptomic differences between WT and *Gata2*^*+/-*^ HSPCs, we performed network analysis using Cytoscape software (Figure 1E). In this analysis, dots represent gene-sets, lines connect gene-sets sharing the same genes and gene-sets that are associated with the same biological processes form clusters indicated by the bigger circles. Earlier studies showed that HSPCs are highly proliferative during EHT *(Zape et al., 2017)*. Strikingly, many gene-sets related to *Cell cycle, DNA repair, Transcription* and *Translation* networks were downregulated in *Gata2*^*+/-*^ HSPCs, indicating that these processes are abrogated (Figure 1E). Conversely, gene-sets related to *Cell junction organization, Integrin signaling, Notch signaling* and *Extracellular matrix formation* were significantly upregulated in *Gata2*^*+/-*^ HSPCs (Figure 1E).

These results suggest that the inability of some embryonic *Gata2*^*+/-*^ HSPCs to complete EHT is due to an defective switch from endothelial programming to hematopoietic programming, possibly preventing their maturation into HSCs.

### Endothelial-repressor transcription factor Gfi1b is downregulated in E11 Gata2^+/-^ HSPCs

Because GSEA and network analysis both indicated that endothelial and hematopoietic genes are enriched in *Gata2*^*+/-*^ HSPCs, the expression of genes defining these transcriptomic signatures was investigated. Endothelial-specific genes such as *Kdr, Cxcl12, Sox17* and *Ramp2* were significantly upregulated in *Gata2*^*+/-*^ HSPCs (Figure 1F), along with an aberrant hematopoietic-specific transcriptome. While some genes indicating hematopoietic programming such as *Notch1* and its targets *Hey1* and *Hes1, Cdkn1c* and *Igf2* were upregulated, other hematopoietic-specific genes such as *Myb* and *Gpr56* (*Adgrg1*) were significantly downregulated in *Gata2*^*+/-*^ HSPCs (Figure 1F). These results support the hypothesis that *Gata2*^*+/-*^ HSPCs can initiate hematopoietic programming but cannot fully gain hematopoietic characteristics due to impaired switching from the endothelial-specific to the hematopoietic-specific transcriptional program. Notably, *Gfi1b* expression was significantly reduced in *Gata2*^*+/-*^ HSPCs (Figure 1F). *Gfi1b* is known to be responsible for the loss of endothelial identity during EHT and its expression is essential for the formation of IAHCs from HECs *(Thambyrajah et al., 2016; Lancrin et al., 2012). Gfi1b* is directly activated by *Gata2* through the hematopoietic specific +16 and +17 kb distal enhancer regions at E11.5 mouse embryos *(Moignard et al., 2013)*. To further investigate whether *Gata2* haploinsufficiency causes *Gfi1b* downregulation due to the reduced activity in the enhancer regions of *Gfi1b*, we sorted CD31^+^cKit^+^ HSPCs from E11 WT and *Gata2*^*+/-*^ AGMs and performed ATAC-sequencing (ATAC-seq) to determine chromatin accessibility. Strikingly, both +16 and +17 kb distal enhancer regions of *Gfi1b* were less accessible in *Gata2*^*+/-*^ HSPCs compared to WT HSPCs indicating *Gata2* regulates *Gfi1b* expression through these distal enhancer regions during EHT (Figure 1G).

### Gata2 haploinsufficiency impairs HSPC maturation during EHT

The effects of *Gfi1b* and incomplete suppression of endothelial identity, as well as the multistep maturational process in *Gata2*^*+/-*^ HSPCs was examined. Previous research showed that HSPCs develop within IAHCs through pro-HSC (*CD31*^+^*cKit*^+^*CD41*^lo^*CD43*^-^*CD45*^-^) → pre-HSC Type I (pre-HSC1 or *CD31*^+^*cKit*^+^*CD41*^lo^*CD43*^+^*CD45*^-^) → pre-HSC Type II (pre-HSC2) and HSC (*CD31*^+^*cKit*^+^*CD41*^lo^*CD43*^+^*CD45*^+^) transitions *(Rybtsov et al., 2014; 2011; Taoudi et al., 2008)*. In this maturation hierarchy, only the most mature compartment (pre-HSC2/HSCs) contains transplantable HSCs and can produce hematopoietic colonies in colony-forming unit culture (CFU-C) *(Taoudi et al., 2008)*.

Because HSC maturation requires activation of *Gfi1b* and consequential downregulation of endothelial genes *(Thambyrajah et al., 2016; Lancrin et al., 2012)*, we asked whether *Gata2* haploinsufficiency causes a block in a specific stage of HSC maturation during EHT. To test this, we dissected WT and *Gata2*^*+/-*^ AGMs from E11 embryos and performed flow cytometry experiments. Using antibody combinations for C*D31, cKit, CD41, CD43* and *CD45* (Figure S1A), the number of pro-HSC, pre-HSC1 and pre-HSC2/HSC cells per in E11 WT and *Gata2*^*+/-*^ AGMs was analysed (Figure 2A). We found that the numbers of E11 pro-HSCs were comparable between WT and *Gata2*^*+/-*^ AGMs indicating the first step of HSPC formation was not hampered in *Gata2*^*+/-*^ embryos (Figure 2B). In contrast, the numbers of pre-HSC1 cells were moderately and pre-HSC2/HSC cells more prominently reduced in E11 *Gata2*^*+/-*^ embryos, suggesting that *Gata2*^*+/-*^ HSPCs cannot complete HSC maturation (Figure 2B).

**Figure 2.**
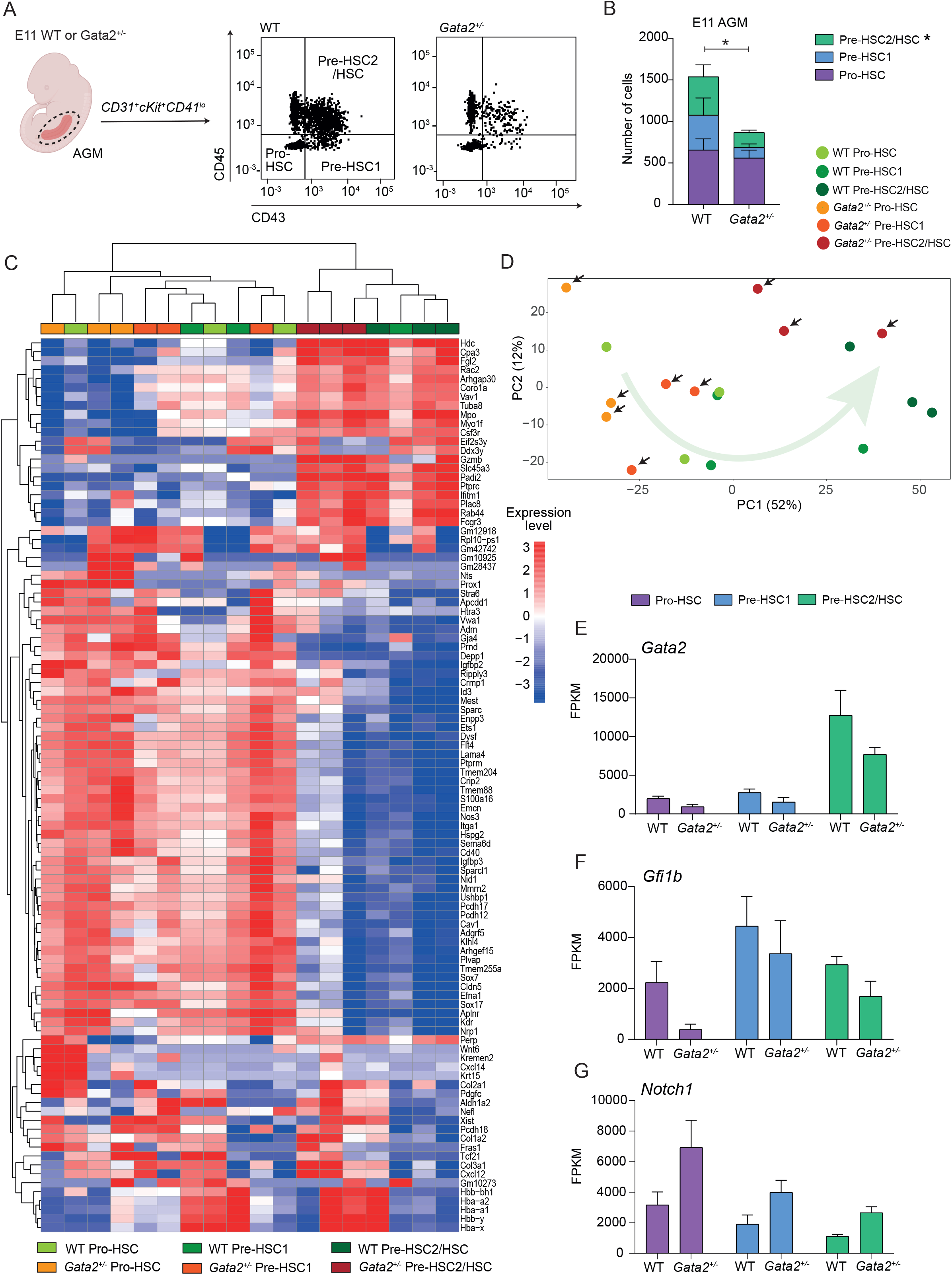
HSPC maturation within IAHCs is impaired in *Gata2*^*+/-*^ during EHT. A) Gating strategy to determine HSPC maturation at E11 WT or *Gata2*^*+/-*^ AGMs. Representative image of pro-HSC, pre-HSC1 and pre-HSC2 gating obtained from WT (left) or *Gata2*^*+/-*^ (right) AGMs. B) Quantification of the number of pro-HSC, pre-HSC1 and pre-HSC2 populations in E11 WT (n=11) or *Gata2*^*+/-*^ (n=13) AGMs. C) Unbiased heatmap of the transcriptomic signatures of pro-HSC, pre-HSC1 and pre-HSC2 populations from 3 independent E11 WT or 3 independent *Gata2*^*+/-*^ AGMs. D) PCA showing the transcriptome of each sample obtained from RNA sequencing. Black arrows indicate *Gata2*^*+/-*^ samples. Green arrow indicates the maturation trajectory based on the transcriptome of WT HSPCs. FPKM values of E) *Gata2* F) *Gfi1b* and G) *Notch1* depicted for each stage of the maturation and compared between WT and *Gata2*^*+/-*^. Error bars represent standard error of the mean. Color code for samples according to the genotype: WT samples in the shades of green and *Gata2*^*+/-*^ samples in the shades of orange. Color code for maturation steps: pro-HSCs, purple; pre-HSC1, blue; pre-HSC2, green. *P < 0.05

To investigate if the maturation of pro-HSCs into pre-HSCs is impaired or delayed in *Gata2*^*+/-*^ AGMs, the flow cytometry analysis included E12 and E13 AGMs dissected from WT and *Gata2*^*+/-*^ embryos. At E12, the numbers of pre-HSC2/HSCs were higher compared to E11 in both WT and *Gata2*^*+/-*^ indicating HSPCs were still actively undergoing maturation at this stage (Figure S1B, comparing to Figure 2B). Both pre-HSC1 and pre-HSC2 cells were significantly reduced in *Gata2*^*+/-*^ compared to WT AGMs at E12 (Figure S1B). The numbers of pro-HSC, pre-HSC1 and pre-HSC2/HSC cells were markedly reduced in both WT and *Gata2*^*+/-*^ AGMs at E13 compared to E11 or E12 (Figure S1B and C, comparing to Figure 2B), however *Gata2*^*+/-*^ AGMs still contained fewer HSPCs at E13 compared to WT AGMs (Figure S1C).

Next, we asked whether the functionality of *Gata2*^*+/-*^ HSCs is altered at E11. Earlier CFU-C studies showed that *Gata2*^*+/-*^ HSPCs produce fewer hematopoietic colonies than WT *(de Pater et al., 2013)*, but it remained unclear whether this was due to a reduction in number of pre-HSC2/HSCs or due to a reduction in their potential to generate CFUs. To address this, we sorted pro-HSC, pre-HSC1 and pre-HSC2/HSC populations from E11 WT and *Gata2*^*+/-*^ AGMs and performed the CFU-C (day 11) assay to quantitatively assess subset hematopoietic potential. Colony counts normalized to the number of cells plated per dish revealed that the functionality of pre-HSC2/HSCs was preserved between WT and *Gata2*^*+/-*^ AGMs, as this population from both genotypes produced similar type and number of hematopoietic colonies (Figure S1D). In addition, pro-HSC and pre-HSC1 populations of both WT and *Gata2*^*+/-*^ AGMs did not form any colonies after 11 days in culture, confirming these cells have not yet acquired HSC potential (*data not shown*).

These results indicate that *Gata2*^*+/-*^ HSPCs are arrested in pro-HSC to pre-HSC maturation during EHT. The ability of *Gata2*^*+/-*^ AGMs to still produce a few pre-HSC2/HSCs indicates that HSC generation is not fully blocked but allows at least some haploinsufficent cells to maintain similar CFU potential as WT Pre-HSC2/HSCs.

### Pre-HSC2/HSCs are marked by unique transcriptomic signatures

To investigate the molecular basis for the *Gata2*^*+/-*^ effects in the pro-HSC stage of maturation, RNA sequencing was performed on pro-HSC, pre-HSC1 and pre-HSC2/HSC compartments from E11 WT and *Gata2*^*+/-*^ AGMs. Differentially expressed genes between WT and *Gata2*^*+/-*^ HSPC subtypes is shown in supplementary material 1-3. Heatmaps comparing the transcriptomic signatures of each cell population from WT and *Gata2*^*+/-*^ AGMs show that overall pro-HSC and pre-HSC1 compartments of both WT and *Gata2*^*+/-*^ have comparable transcriptomes which dramatically change during pre-HSC2/HSC maturation (Figure 2C). PCA confirmed that, in both WT and *Gata2*^*+/-*^, the transcriptome of pre-HSC2/HSCs cluster separately from the pro-HSC and pre-HSC1 compartments (Figure 2D). Pre-HSC2/HSCs from both genotypes uniquely express *Ptprc* (*CD45*), are marked by the specific expression of *Hdc, Fgl2, Slc45a, Padi2, Rab44* and *Fcgr3* (Figure 2C), and *Flt4, Ptprm, Itga1* and *Sox17* expression was extinguished in pre-HSC2/HSCs of both WT and *Gata2*^*+/-*^ to indicate that mature HSCs acquire a unique transcriptomic signature (Figure 2C).

### Gata2^+/-^ HSPCs fail to repress endothelial programming throughout HSC maturation

Flow cytometry analysis showed that HSPC maturation in *Gata2*^*+/-*^ was predominantly impaired during Pro-HSC to Pre-HSC1 transition (Figure 2B, Figure S1B). Comparison of the transcriptomes of WT and *Gata2*^*+/-*^ by heatmap analysis and PCA, showed that the transcriptome profiles of WT pre-HSC1 cells were distributed between pro-HSC and pre-HSC2/HSCs states. However, *Gata2*^*+/-*^ pre-HSC1 cells were transcriptionally closer to a pro-HSC-like state (Figure 2C and D). Although some endothelial-specific genes like *Sox7* and *Flt4* were expressed in both WT and *Gata2*^*+/-*^ pre-HSC1 cells (Figure 2C), other endothelial-specific genes such as *Igfbp2, Vwa1* and *Gja4* were upregulated in *Gata2*^*+/-*^ compared to WT pre-HSC1 cells (Figure S2A-C and supplementary material 2).

WT and *Gata2*^*+/-*^ pre-HSC2/HSCs did not show striking differences in transcriptome (Figure 2C and supplementary material 3) probably explaining the preserved functionality in CFU-C assays. Expression of some endothelial-specific genes such as *Sox17, Kdr* and *Flt1* were silenced during pre-HSC1 to pre-HSC2/HSC maturation in both WT and *Gata2*^*+/-*^ cells (Figure 2C), while others such as *Cxcl12, Col3a1* and *Aplnr* were upregulated in *Gata2*^*+/-*^ pre-HSC2/HSCs compared to WT. (Figure 2C, Figure S2D-F).

These differences indicate that the mature Pre-HSC1 and Pre-HSC2/HSC compartments of *Gata2*^*+/-*^ are transcriptionally distinguishable from WT and suggest that *Gata2*^*+/-*^ HSPCs incompletely repress endothelial programming through maturation from pro-HSC to pre-HSC1 and pre-HSC2/HSC. However, some *Gata2*^*+/-*^ HSPCs are still capable of undergoing complete AGM maturation despite carrying endothelial transcriptomic signatures.

### Gata2^+/-^ HSPCs incompletely activate hematopoietic programming throughout HSC maturation

Because some hematopoietic transcriptomic signatures were upregulated whereas others were downregulated in *Gata2*^*+/-*^ HSPCs (Figure 1F), we asked whether this observation was due to the reduced number of mature pre-HSC2/HSCs within HSPCs. To explore this, the expression levels of hematopoietic-specific *cKit, CD44* and *Myb* genes were examined at the individual steps of maturation. For both WT and *Gata2*^*+/-*^, expression of these genes increased throughout the maturation (Figure S2G-I). However, levels were reduced in *Gata2*^*+/-*^ HSPCs compared to WT in all maturation stages, confirming that hematopoietic transcriptional programming is hampered in all *Gata2*^*+/-*^ HSPCs (Figure S2G-I). Strikingly, the most significant reduction was in the *Gata2*^*+/-*^ pro-HSCs (Figure S2G-I and supplementary material 1).

Hematopoietic specific genes such as *Vav1, Rac2* and *Mpo* were expressed in WT HSPCs at all stages and levels gradually increased throughout maturation (Figure S2J-L). Expression of these genes was downregulated most prominently in pro-HSCs and only activated later during the pre-HSC1 and pre-HSC2/HSCs maturation in *Gata2*^*+/-*^ cells (Figure S2J-L), indicating a direct effect of Gata2 haploinsufficiency in the onset of the hematopoietic programming.

Hence *Gata2* likely plays a crucial role in HSPCs undergoing EHT to downregulate endothelial identity and upregulate hematopoietic transcriptional programming to promote a complete transition to the HSC-like state.

### Gfi1 and Gfi1b are downregulated in Gata2^+/-^ HSPCs during pro-HSC to pre-HSC maturation

Previous studies showed that *Gata2* is upregulated in HSPCs compared to ECs and HECs during EHT *(Eich et al., 2018)*. By comparing the expression level of *Gata2* among pro-HSC, pre-HSC1 and pre-HSC2/HSC compartments we found that *Gata2* expression levels are highest in the most mature compartment (pre-HSC2/HSC) of HSPCs (Figure 2E) and confirmed that *Gata2*^*+/-*^ HSPCs in all maturation stages have reduced *Gata2* expression compared to WT (Figure 2E).

We next examined whether the expression of *Gfi1b*, a known repressor of endothelial gene expression, is altered in *Gata2*^*+/-*^ embryos. *Gfi1b* expression levels were reduced in *Gata2*^*+/-*^ Pro-HSCs, while pre-HSC1 and pre-HSC2/HSC compartments showed only a marginal reduction in *Gfi1b* expression level compared to WT (Figure 2F). These results suggest that *Gata2* haploinsufficiency impairs *Gfi1b* activation predominantly in the pro-HSC stage of the HSPC maturation.

Previous studies showed that *Gfi1*, a highly homologous interaction partner of *Gfi1b*, is also required for IAHC formation during EHT *(Thambyrajah et al., 2016; Lancrin et al., 2012). CD45*^+^ pre-HSC2/HSCs co-express *Gfi1b* and *Gfi1*, indicating that their co-activation is required during HSPC maturation within IAHCs *(Thambyrajah et al., 2016)*. Therefore, we examined whether *Gfi1* expression was altered in *Gata2*^*+/-*^ HSPCs throughout maturation. Strikingly, *Gfi1* expression was completely abolished in *Gata2*^*+/-*^ pre-HSC1 cells (Figure S2M). This suggests *Gata2* is required for the expression of both *Gfi1* genes that are crucial for the repression of endothelial programming during HSPC maturation.

### Notch signaling is upregulated in Gata2^+/-^ HSPCs

*Notch signaling* is essential for EHT and HSPC maturation and *Notch1* and its targets such as *Hes1* are also required for the initiation of EHT in the AGM *(Souilhol et al., 2016; Robert-Moreno et al., 2005)*. However, AGM HSCs become *Notch*-independent throughout maturation, with continued high *Notch1* activity eventually blocking HSC maturation *(Souilhol et al., 2016)*. Because our E11 gene-set analysis showed upregulated *Notch signaling Gata2*^*+/-*^ HSPCs, we explored the expression levels of individual key *Notch signaling* mediators. Both *Notch1* and *Hes1* were enriched in *Gata2*^*+/-*^ HSPCs compared to WT (Figure 2G and S2N) whereas the expression level of *Notch2* was comparable between WT and *Gata2*^*+/-*^ HSPCs throughout maturation (Figure S2O). In accordance with *Gata2* being a known downstream target of *Notch1*, these results suggest a possible feedback regulation of *Notch signaling* by *Gata2* during EHT and HSPC maturation.

### Both IAHCs and single bulging cells are diminished in Gata2^+/-^ AGMs

Our results indicated that HSPC maturation within IAHCs is inhibited in *Gata2*^*+/-*^ AGMs. A recent study reported that all IAHCs express Gata2 but that *Gata2*-expressing CD27^+^ single bulging cells (SBC, or sIAHC) in 1-to 2-cell clusters are HSCs *(Vink et al., 2020)*. To clarify whether *Gata2* haploinsufficiency exclusively diminishes HSPC maturation within IAHCs or has an effect on these SBCs, we dissected AGMs from E11 WT and *Gata2*^*+/-*^ embryos and performed whole-mount immunofluorescence staining for CD31 and cKit (Figure 3A-C). In whole WT and *Gata2*^*+/-*^ AGMs, we quantified the number of CD31^+^cKit^+^ SBCs and the CD31^+^cKit^+^ cells located in IAHCs (Figure 3D; IAHC, E; SBC or sIAHC and G). We also assessed the number of single CD31^+^cKit^+^ cells that are not attached to endothelium or IAHCs and are found in the aortic lumen (AL) (Figure 3F and G). Our results showed that the number of CD31^+^cKit^+^ cells within all AGM compartments are decreased in E11 *Gata2*^*+/-*^ embryos, with significant reductions in both IAHCs and SBCs (Figure 3G). Furthermore, because *CD27* is expressed in both SBCs and some IAHCs, and is also a marker for multipotent HSPCs *(Vink et al., 2020)*, we examined the expression level of CD27 throughout the maturation stages. CD27 expression levels were greatly reduced in *Gata2*^*+/-*^ pre-HSC1 compartment (Figure 3H) to suggest that *Gata2* haploinsufficiency does not exclusively affect the HSC maturation within IAHCs. It additionally reduces the HSC potential within AGM HSPC pool, thus indicating a broader function for *Gata2* during EHT.

**Figure 3.**
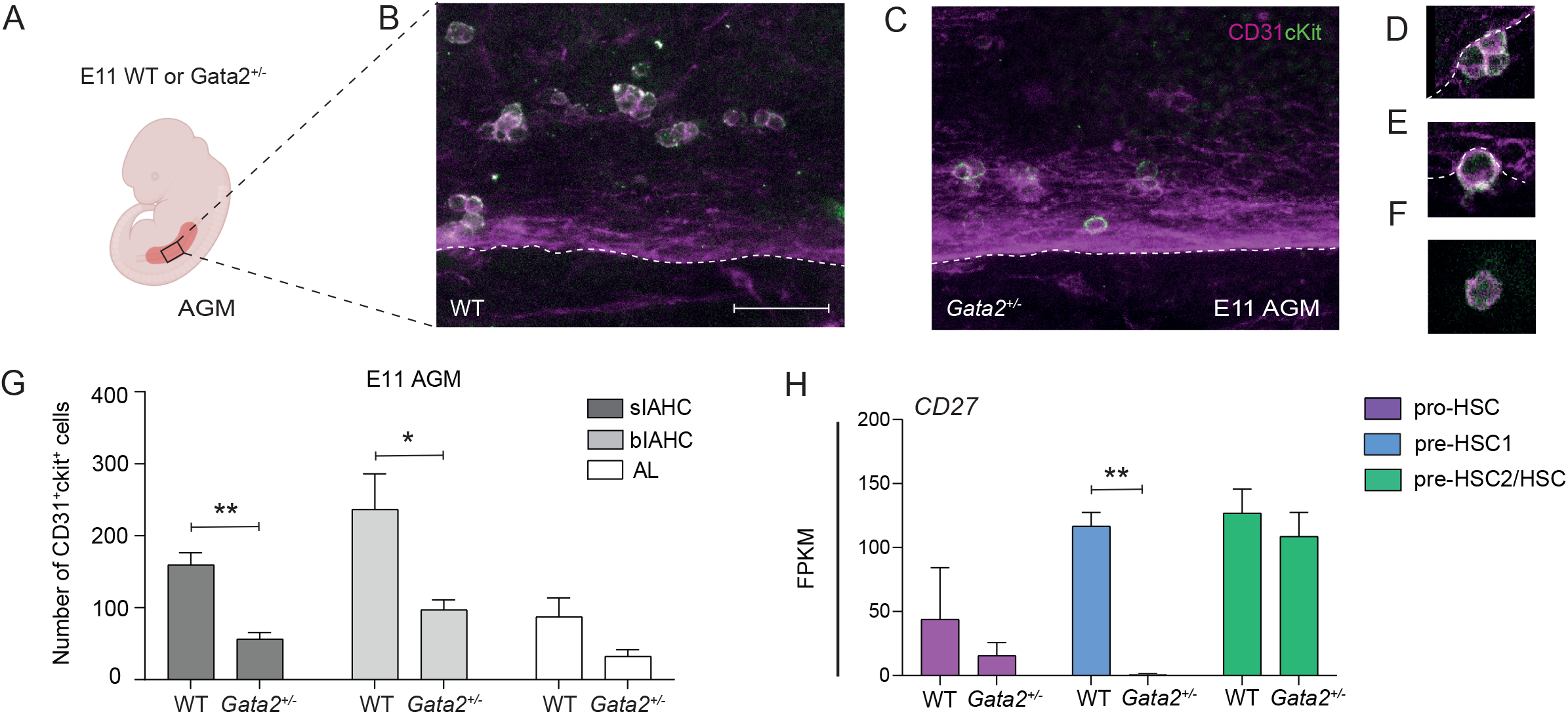
Both IAHCs and single bulging cells are depleted in *Gata2*^*+/-*^ AGMs. A) Illustration of AGM region dissected for analysis from E11 WT or *Gata2*^*+/-*^ embryos. B-F) Representative images of CD31^+^cKit^+^ cells obtained by confocal imaging at E11 B) WT AGM, C) *Gata2*^*+/-*^ AGM, D) IAHCs, E) SBC or sIAHC and F) AL cell. White bar indicates 50 um in length. G) Quantification of CD31^+^cKit^+^ cells located in IAHCs, as SBCs and as AL cell within E11 WT (n=3) and *Gata2*^*+/-*^ (n=4) AGMs. H) FPKM values of CD27 depicted for each stage of the maturation and compared between WT and *Gata2*^*+/-*^. Error bars represent standard error of the mean. *P < 0.05, **P < 0.01. AGM, aorta-gonad-mesonephros; IAHC, intra-aortic hematopoietic cluster; SBC, single bulging cell; AL, aortic lumen cell.

### Hematopoietic-specific gfi1b induction restores embryonic HSCs in gata2b^-/-^ zebrafish

To test whether the effect of *Gata2* haploinsufficiency on EHT can be overcome by inducing *Gfi1b* expression, we took advantage of the previously described *gata2b*^*-/-*^ zebrafish model *(Gioacchino et al., 2021)*. Definitive HSPCs in *gata2b*^*-/-*^ zebrafish are marked by reduced *cmyb* signal from 33 hours post fertilization (hpf) and onwards and the number of HSCs (*CD41*^*int*^) are severely depleted in *gata2b*^*-/-*^ embryos at 3 days post-fertilization (dpf) *(Gioacchino et al., 2021)*. To test the effect of ectopic *gfi1b* expression on *gata2b*^*-/-*^ HSPCs, we injected a HEC tissue specific *gfi1b* expression cassette into WT and *gata2b*^*-/-*^ *CD41:GFP* embryos at 1-cell stage (Figure 4A). We found that, upon *gfi1b* induction, *cmyb* signals were normalized in *gata2b*^*-/-*^ HSPCs at 33 hpf (Figure 4B). In addition, the number of *CD41*^*int*^ HSCs in *gata2b*^*-/-*^ zebrafish was restored to WT levels at 3 dpf (Figure 4C-E).

**Figure 4.**
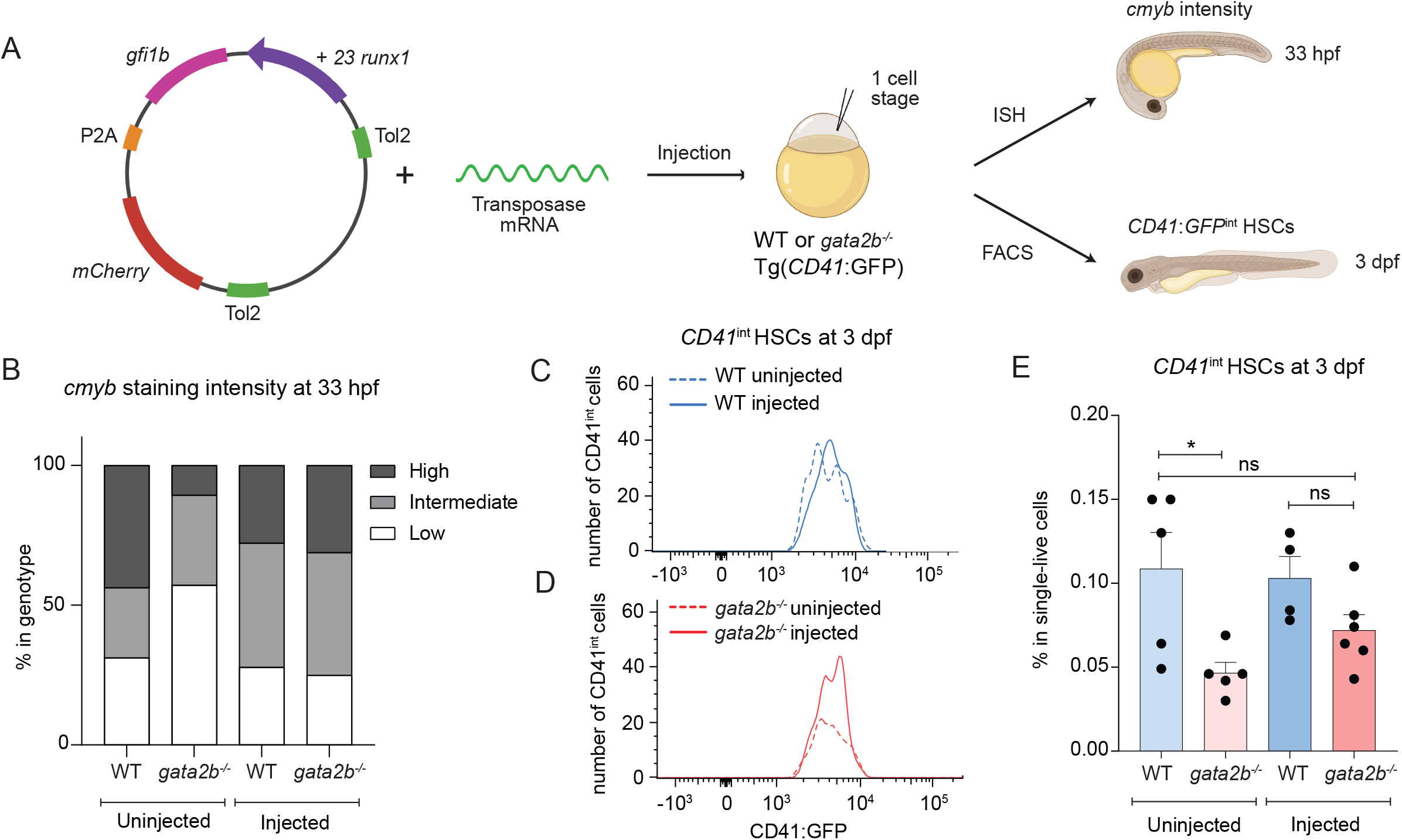
*gfi1b* induction restores the number of embryonic HSCs in *gata2b*^*-/-*^ zebrafish. A) Illustration of the experiment and analysis strategy for *gfi1b* induction in zebrafish embryos. Rescue construct containing +23 *runx1* enhancer, *gfi1b* and *mCherry* is injected to the single cells of Tg(CD41:GFP) WT or *gata2b*^*-/-*^ zebrafish embryos in combination with transposase mRNA. Embryos were analyzed at 33 hpf for *cmyb* signal intensity and at 3 dpf for CD41^+^ HSCs. B) cmyb signal intensity analyzed at 33 hpf by ISH and compared between uninjected (WT n=2, *gata2b*^*-/-*^ n=16) and injected (WT n=18, *gata2b*^*-/-*^ n=16) WT and *gata2b*^*-/-*^ embryos. C) Representative image of the number of CD41^int^ HSCs compared between uninjected (n=5) and injected (n=4) groups of WT and D) uninjected (n=5) and injected (n=6) groups of *gata2b*^*-/-*^ embryos. E) Proportion of CD41^int^ HSCs compared between uninjected and injected groups of WT and *gata2b*^*-/-*^ embryos. Dots represent individual samples and each sample contains 4 pooled embryos. ISH, in situ hybridization; FACS, fluorescent activated cell sorting.

These results show that ectopic *gfi1b* expression can rescue the embryonic phenotype of *gata2b*^*-/-*^ zebrafish and indicate that *Gata2* regulates Gfi1b during EHT.

## Discussion

In this study, we aimed to assess the role of *Gata2* on the generation of phenotypic HSCs through EHT. Although HSPCs do not completely switch-off their endothelial gene expression and remain expressing endothelial markers like *CD31* throughout EHT and maturation, moderated repression of endothelial programming is essential for the maturation and thus for establishing the HSC fate *(Oatley et al., 2020; Baron et al., 2018; Zhou et al., 2016; Swiers et al., 2013)*. We showed that *Gata2* haploinsufficiency does not completely abrogate the hematopoietic programming during EHT, but reduces the ability of HSPCs to complete HSC maturation in the AGM.

*Gata2*^*+/-*^ HSPCs were predominantly blocked during pro-HSC to pre-HSC maturation and functional HSCs (pre-HSC2/HSCs) were significantly reduced at both E11 and E12 AGMs. We showed that phenotypic *Gata2*^*+/-*^ HSCs (pre-HSC2/HSCs) not only fail to repress endothelial genes like *Cxcl12* and *Col3a1*, they also incompletely activate hematopoietic genes like *cKit* and *Myb* during AGM maturation. Together this implies that *Gata2* acts as a mediator in the gene regulation of the endothelial and hematopoietic transcriptional programs during EHT. Previous studies showed that *Gata2*^*-/-*^ embryos do not form IAHCs in the AGM implying that *Gata2* expression is required for EHT *(de Pater et al., 2013; Tsai et al., 1994)*. Our results extend these observations by showing that *Gata2* has multiple roles during EHT and is a critical novel regulator of HSPC maturation.

Despite the fact that HSC numbers were reduced in *Gata2*^*+/-*^ AGMs and their transcriptome was distinct from WT HSCs, CFU assays showed that their functionality was preserved. These results revise the observations from previous studies where it was shown that colony forming ability of *Gata2*^*+/-*^ AGM HSPCs is reduced *(de Pater et al., 2013)* and now clarify that this is due to impaired HSC maturation in *Gata2*^*+/-*^ AGMs and not to a reduction in HSC functionality. However, CFU assays are limited when testing repopulating ability and lymphoid potential of HSCs and therefore transplantation studies are needed to elucidate the true potential of embryonic *Gata2*^*+/-*^ HSCs. It is expected that these cells will be less robust when transplanted competitively with WT AGM HSCs, as has been previously shown for BM HSCs (*Ling et al., 2004*). *Gfi1b* was downregulated and hematopoietic distal enhancers of *Gfi1b* were less active in *Gata2*^*+/-*^ HSPCs. Because *Gfi1b* expression is required to repress endothelial programming in HSPCs and for the formation of IAHCs during EHT, we propose that *Gata2* activates *Gfi1b* through its +16 and +17 distal enhancer regions to repress the endothelial identity of HSPCs. Restored number of HSCs in *gata2b*^*-/-*^ zebrafish embryos upon ectopic *gfi1b* expression validated that *Gata2* activates *Gfi1b* and this regulatory mechanism is essential for the generation of embryonic HSCs. Although previous studies provided in silico and experimental evidence suggesting *Gata2* and *Gfi1b* regulate each other *(Moignard et al., 2013)*, to our knowledge this study is the first experimental proof pointing out the phenotypic consequences of the disruption of this regulatory mechanism during EHT.

Importantly, we found an increased activity of *Notch signaling* in *Gata2*^*+/-*^ HSPCs. Previous studies in mice and zebrafish showed that *Notch signaling* is an upstream activator of *Gata2 (Dobrzycki et al., 2020; Guiu et al., 2012)*. Because the expression of *Gata2* is reduced in *Gata2*^*+/-*^ HSPCs, increased *Notch signaling* could be activated as a compensatory mechanism in these cells. Therefore, our results suggest a possible feedback regulation of *Notch signaling* through *Gata2* expression. Because it was shown that downregulation of *Notch signaling* is required during HSPC maturation (*Souilhol et al., 2016*), we cannot exclude the possibility of impaired HSPC maturation in *Gata2*^*+/-*^ AGMs is influenced by the high *Notch* activity. Although inducing *gfi1b* can sufficiently rescue the embryonic phenotype in *gata2b*^*-/-*^ embryos, the contribution of *Notch signaling* herein requires further investigation.

Finally, we showed that many gene-sets related to *Cell cycle, Proliferation* and *DNA repair* were downregulated in *Gata2*^*+/-*^ HSPCs throughout all maturation stages (Figure S3A-C). Although decreased proliferative signatures are hallmarks of impaired HSPC maturation, whether this has an influence on genome stability of *Gata2*^*+/-*^ HSPCs remains to be explored. Because HSCs generated through EHT during embryonic stages are the source of the adult HSC pool, further studies are needed to understand the influence of prenatal *GATA2* haploinsufficiency to HSC fitness after birth and throughout adulthood.

## Acknowledgements

We thank members of the de Pater lab for helpful discussions. We thank the Experimental Animal Facility of Erasmus MC for animal husbandry and the Erasmus Optical Imaging Center for confocal microscopy services. This research is supported by the European Hematology Association (junior non clinical research fellowship)(EdP), the Dutch Cancer Foundation KWF/Alpe d’HuZes (SK10321)(EdP) and the Daniel den Hoed Foundation for support of the Cancer Genome Editing Center (IT).

## Authorship contributions

EdP and CK conceived the study; CK, EG, DV, HdL, JZ, MH, and EB performed experiments; CK, EG, DV, RH and EdP analysed results; IT provided resources and CK and EdP wrote the manuscript and IT and ED revised the manuscript.

## Disclosures

The authors declare no conflicts of interests

## Notes

### Competing Interest Statement

The authors have declared no competing interest.

