## Supplementary figures and images for "*Gata2*-regulated *Gfi1b* expression controls endothelial programming during endothelial-to-hematopoietic transition"

### supplementary figure 1-3

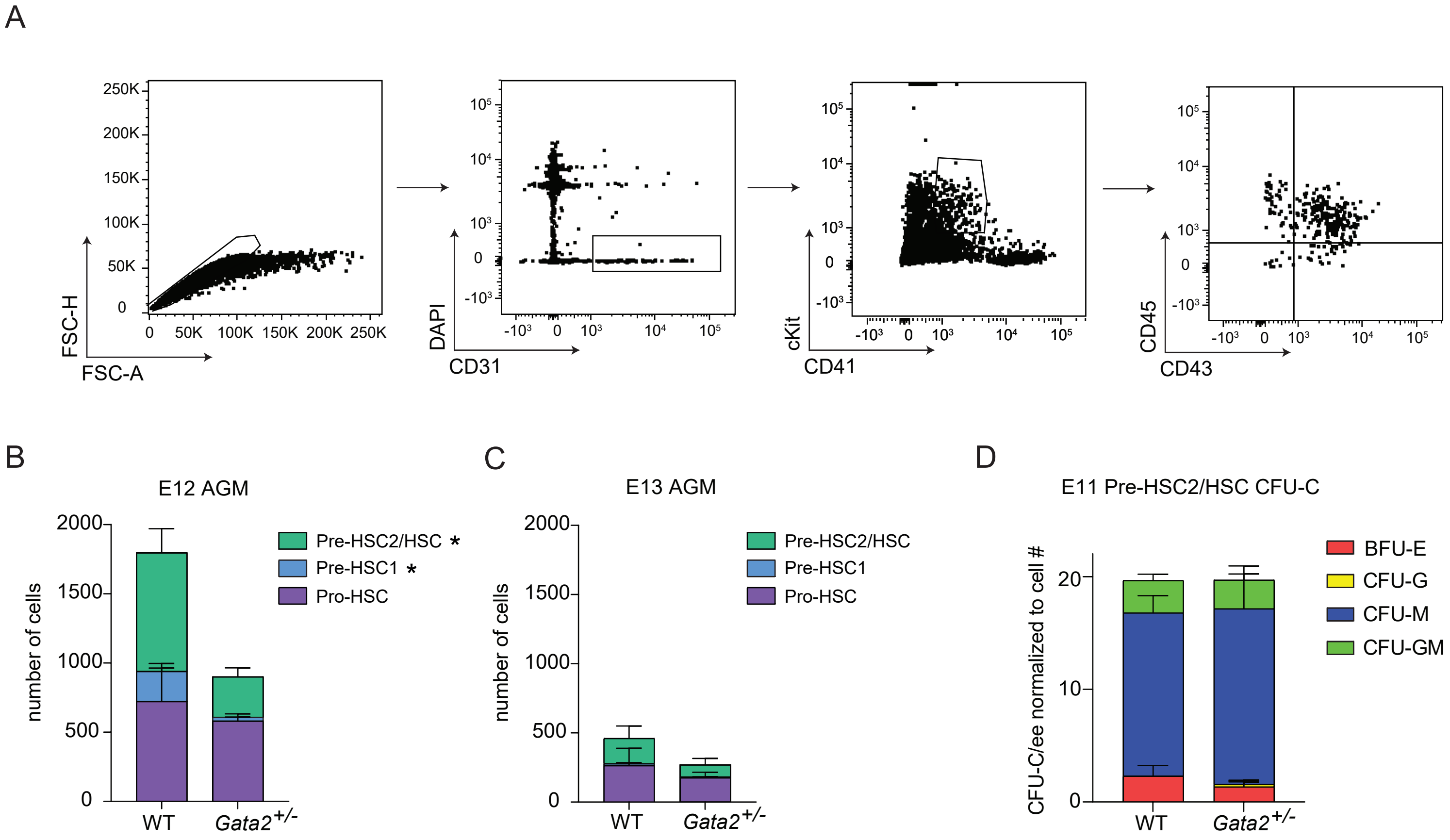


Supplementary figure 1

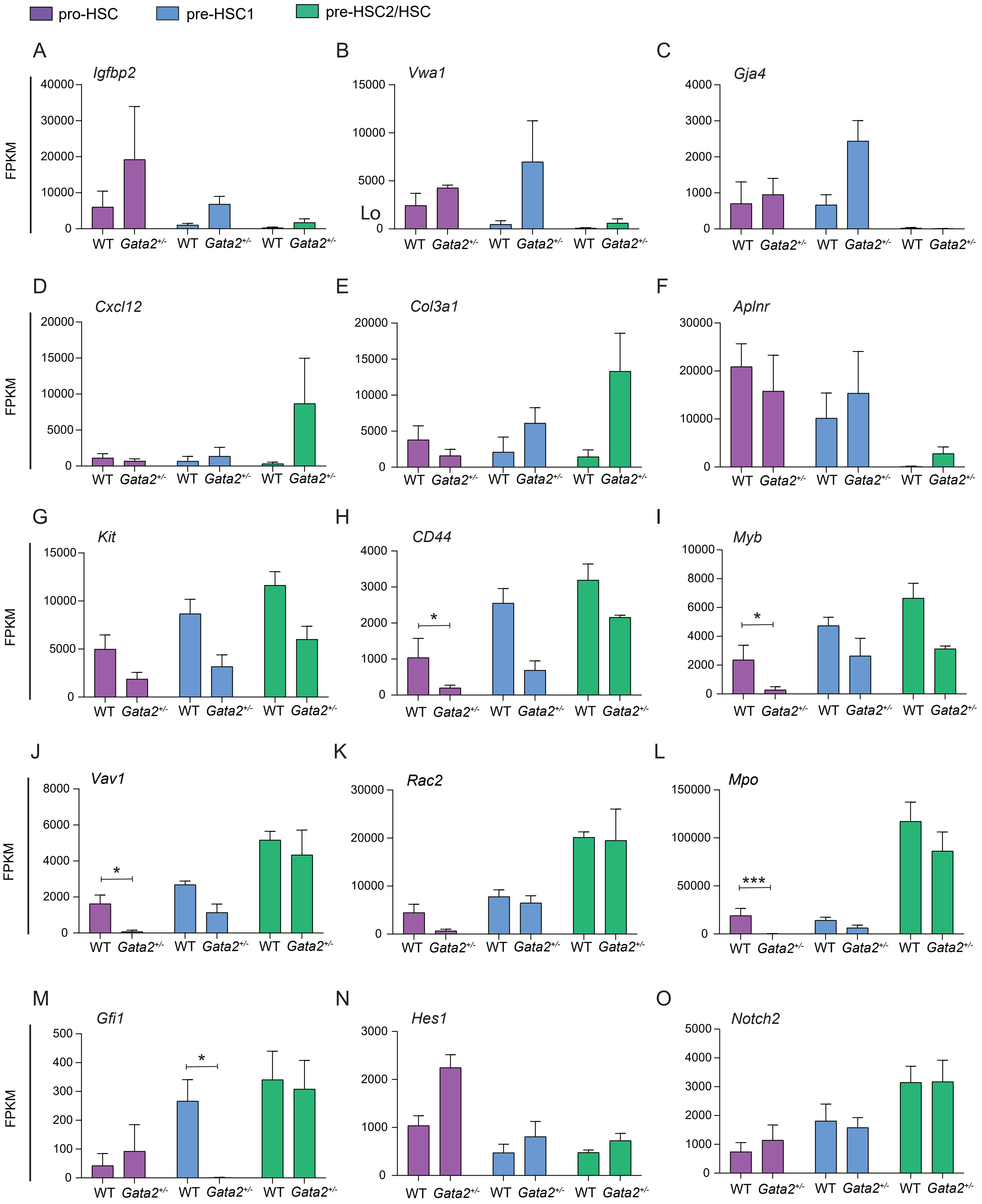


Supplementary figure 2

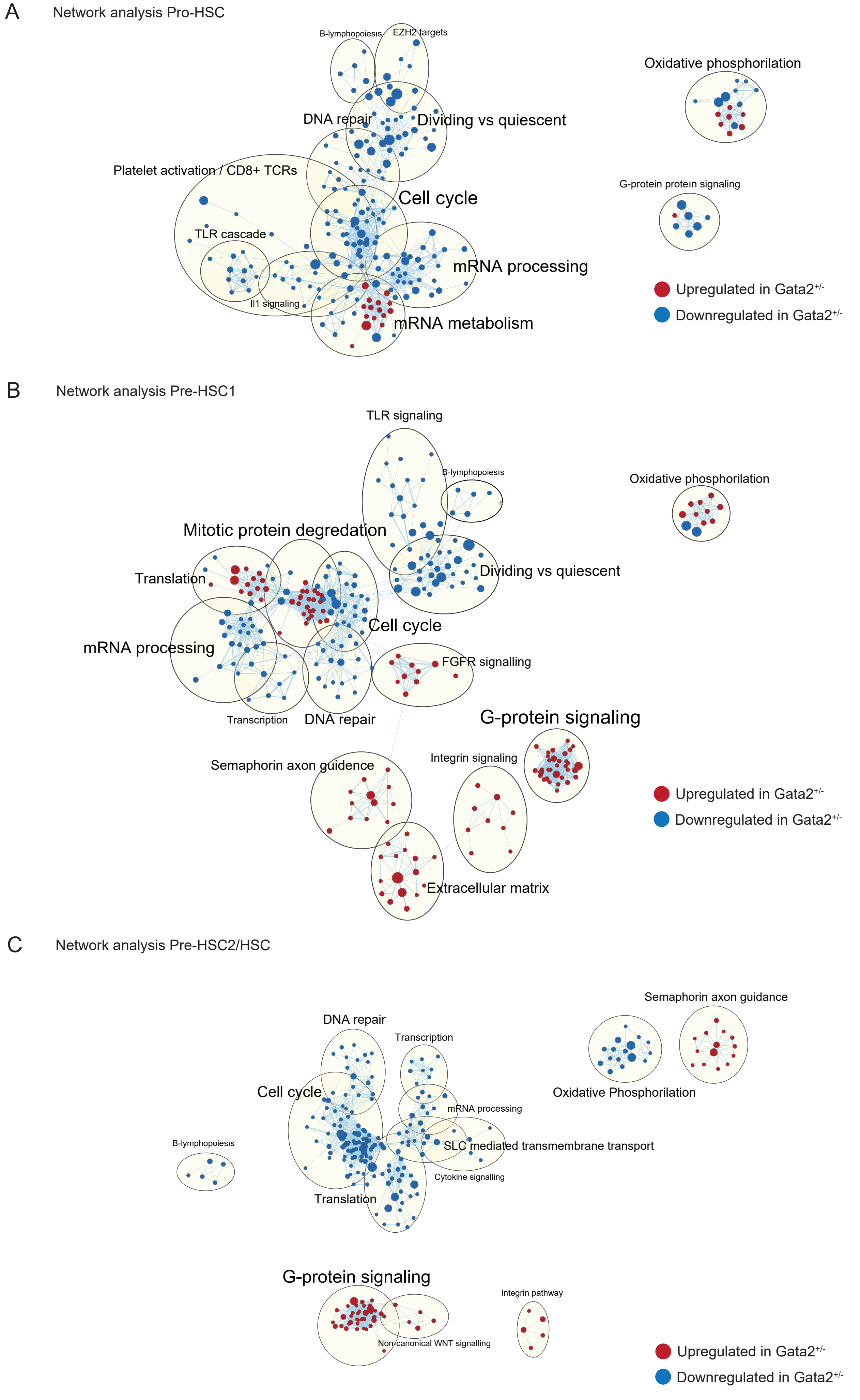


Supplementary figure 3
