## supplementary information for "*Gata2*-regulated *Gfi1b* expression controls endothelial programming during endothelial-to-hematopoietic transition"

**Table S1. Primers used for Gibson assembly reaction.**

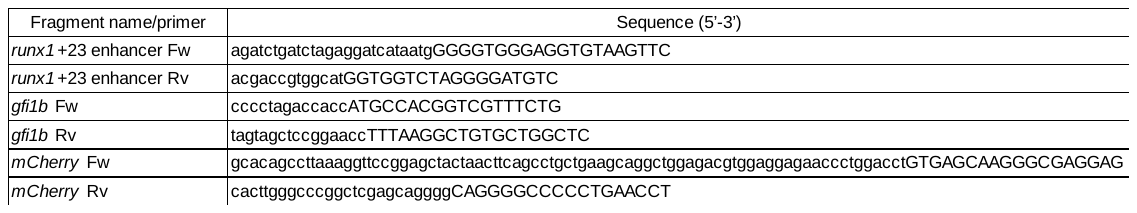

**Table S2. CD31^+^cKit^+^ cell numbers in E11 WT and *Gata2^+/-^* AGMs.**

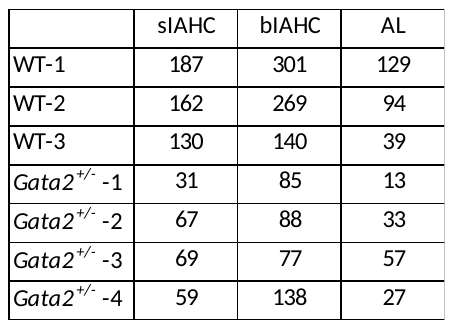

**Supplementary figure 1. HSPC maturation is not delayed but impaired in *Gata2^+/-^* AGMs.**

A) Gating strategy to detect HSPC maturation in AGM. B) Quantification of the number of pro-HSC, pre-HSC1 and pre-HSC2 populations in E12 WT or *Gata2^+/-^* AGMs. C) Quantification of the number of pro-HSC, pre-HSC1 and pre-HSC2 populations in E13 WT or *Gata2^+/-^* AGMs. D) Quantification of BFU-E, CFU-G, CFU-M and CFU-GM colonies after 11 days of CFU culture of E11 WT and *Gata2^+/-^* pre-HSC2/HSCs. Colony numbers were normalized to the number of cells plated per dish.

**Supplementary figure 2. *Gata2* haploinsufficiency impairs both endothelial and hematopoietic transcriptional programming throughout HSPC maturation.**

A) Comparison of FPKM values of *Vav1*, B) *Rac2*, C) *Mpo,* D) *Igfbp2,* E) *Vwa1,* F) *Gja4,* G) *Cxcl12*, H) *Col3a1*, I) *Aplnr,* J) *Gfi1*, K) *Hes1,* L) *Notch2,* M) *Kit,* N) *Cd44* and O) *Myb* between WT or *Gata2^+/-^* pro-HSC, pre-HSC1 and pre-HSC2/HSCs. Color code for HSPC maturation steps: pro-HSCs, purple; pre-HSC1, blue; pre-HSC2, green. *P < 0.05, **P < 0.01, ***P < 0.001

**Supplementary figure 3. Proliferation is abrogated in *Gata2^+/-^* HSPCs throughout maturation.**

A) GSEA network analysis of pro-HSC, B) pre-HSC1 and C) pre-HSC2 populations. WT and *Gata2^+/-^* HSPCs were compared. Red dots are showing the upregulated and blue dots are showing the downregulated gene sets in *Gata2^+/-^* HSPCs compared to WT.

**Supplementary material 1. Differentially expressed genes in *Gata2^+/-^* pro-HSCs compared to WT pro-HSCs.**

**Supplementary material 2. Differentially expressed genes in *Gata2^+/-^* pre-HSC1 cells compared to WT pre-HSC1 cells.**

**Supplementary material 3. Differentially expressed genes in *Gata2^+/-^* pre-HSC2/HSCs compared to WT pre-HSC2/HSCs.**
